# Towards functional annotation with latent protein language model features

**DOI:** 10.1101/2025.10.02.680154

**Authors:** Jake Silberg, Elana Simon, James Zou

**Affiliations:** Stanford University

## Abstract

Protein Language Models (PLMs) create high-dimensional embeddings that can be transformed into interpretable sparse features using Sparse Autoencoders (SAEs), where each feature activates on specific protein elements or patterns. However, scalably identifying which features are cohesive and reliable enough for protein annotation remains challenging. We address this by developing a validation pipeline combining three complementary methods: (1) expanded database matching across 20+ annotation sources including hierarchical codes, (2) feature-guided local structural alignment to identify structurally consistent activation regions, and (3) LLM-based feature description generation. Our annotation pipeline demonstrates three key properties of SAE features that make them a useful source of functional annotation complementary to existing methods. First, they can represent more granular patterns than existing protein databases, enabling the identification of sub-domains. Second, they can detect missing annotations by finding proteins that display recognizable structural motifs but lack corresponding database labels. Here, we automatically identify at least 491 missing CATH topology annotations with our pipeline. Third, they can maintain structural consistency across unseen proteins. Of our 10,240 SAE features, we find 615 that are consistently structurally similar in unannotated metagenomic proteins, allowing us to structurally match at least 8,077 metagenomic proteins to characterized proteins. This provides a rapid annotation pipeline with constant time search regardless of database size, that automatically includes structural and function information about the feature that triggered the match. Code is available at https://github.com/jsilbergDS/towards_functional_annotations_plms

## 1 Introduction

Proteins play essential roles in nearly all biological processes, yet our ability to annotate their functional elements lags behind recent growth in sequence data. With the advent of metagenomics, vast datasets of protein sequences have been identified, but many remain functionally uncharacterized (Karin and Steinegger [2025]). Traditional computational methods annotate proteins based on similarity to well-characterized peers, but these approaches struggle when sequence similarity is low (Karin and Steinegger [2025]).

Recent advances in protein language models (PLMs) like ESM-2 (Lin et al. [2023]) have demonstrated remarkable ability to capture evolutionary patterns across the protein universe. However, ESM embeddings cannot easily be translated into interpretable features that represent specific protein domains. Sparse autoencoders (SAEs) have emerged as powerful tools for mechanistic interpretability. When applied to PLMs, they can decompose dense embeddings into interpretable features that frequently correspond to functional protein elements without biological supervision (Simon and Zou [2024]), (Adams et al. [2025]). This suggests that SAE features could serve as a foundation for efficient functional annotation. However, not all SAE features are suitable for annotation. Some may be polysemantic, activating on multiple unrelated concepts, while others may represent patterns meaningful to the model but not interpretable by humans who rely on structural, sequential, and functional annotations.

To harness SAE features for practical annotation, we need to systematically identify features that correspond to interpretable biological concepts. To identify cohesive features, we focus on those that activate most strongly on groups of proteins that are functionally or structurally related, as defined by meeting one of three validation criteria:

1. A single annotation from an existing annotation database is predictive of feature activation
2. The activating regions of the highest activating proteins are structurally similar
3. A language model can generate a feature description that is predictive of activation, as validated in a held-out set of proteins

In this work, we analyze latent features from an SAE trained on Layer 18 of ESM-2 650M, yielding 10,240 features (referred to throughout as f/feature-number, e.g., f/401). Our analysis proceeds in two main parts. First, we develop and evaluate our three-pronged validation approach, demonstrating how expanded database matching, feature-guided local structural alignment, and LLM-based pattern recognition complement each other to identify cohesive features. We show this approach doubles annotation coverage, identifying over 60% of features with strong correspondence to biological annotations (*F* 1 > 0.8). With local structural alignment, we find an additional 2.4% of the features with worse correspondence to existing databases, but high structural similarity.

Second, we demonstrate three practical applications that highlight the advantages of SAE features for protein annotation: (1) capturing granular subdomains within existing annotations that reveal discrete functional units, (2) detecting missing annotations by identifying proteins with recognizable structural motifs that lack corresponding database labels, and (3) enabling zero-shot generalization to novel metagenomic proteins, including those without matches to existing protein families, specifically, Pfam (Bateman et al. [2004]).

Utilizing PLM SAE features in this annotation pipeline offers natural advantages: First, because the top structures for each feature can be pre-computed, during a search, hits can be found in constant-time search regardless of database size. Second, because we know the feature(s) triggering the hit, the search can automatically include known structural or functional information about that feature (such as its links to existing databases or a natural language description). Finally, because the features are found in an unsupervised manner, the search can find structural hits based on features not in existing databases.

## 2 Related Work

### Traditional Sequence-Based Annotation Methods

Functional annotation of uncharacterized proteins has historically relied on sequence conservation approaches. Hidden Markov Models and Position-Specific Scoring Matrices form the foundation of major databases including CATH (Orengo et al. [1997]) (via Gene3D Buchan et al. [2002]) and Pfam (Bateman et al. [2004]). These have been integrated into unified resources like UniProt Consortium [2015] and InterPro (Hunter et al. [2009]), with search tools like InterProScan enabling annotation of arbitrary sequences (Jones et al. [2014]). However, these methods struggle with divergent sequences, particularly from metagenomic sources, leading to specialized databases like Novel Metagenomic Pfams (NMPfamsDB) (Baltoumas et al. [2024]) that specifically curate sequences lacking Pfam annotations.

### Structure-Based Approaches

AlphaFold 2 (Jumper et al. [2021]) catalyzed a shift toward 3D structure-based annotation methods. This has enabled new databases focused on structural similarity, such as The Encyclopedia of Domains (Lau et al. [2024]) based on the Merizo tool (Lau et al. [2023]). However, large-scale structural searching remains computationally challenging despite algorithmic advances like TMalign (Zhang and Skolnick [2005]) and CEalign (Shindyalov and Bourne [1998]). FoldSeek addresses this by converting structural search into sequence matching using a 3D-informed alphabet, achieving orders-of-magnitude speedup (van Kempen et al. [2022]). Merizo-search uses embeddings trained on CATH domains for rapid structure matching (Kandathil et al. [2025]). While powerful, these approaches have limitations: FoldSeek doesn’t automatically provide domain-specific functional information about the region(s) of each protein that triggered the match, and Merizo-search is constrained to identify hits based on supervised training on CATH annotations.

### Interpretable Features from Protein Language Models

Recent work has explored sparse autoencoders (SAEs) for extracting interpretable features from PLMs. InterPLM (Simon and Zou [2024]) demonstrated correspondence between amino acid activations and UniProtKB annotations, while InterProt (Adams et al. [2025]) associated protein-level activations with Pfam families. However, these approaches achieved limited coverage, leaving over 75% of features unexplained when using stringent matching criteria.

Our work extends this foundation by: (1) incorporating the full InterPro database across 20+ annotation sources including hierarchical codes, (2) developing systematic structural validation through feature-guided local structural alignment, and (3) integrating LLM-based pattern recognition to identify features missed by existing codes. This comprehensive approach doubles annotation coverage while enabling practical applications for both missing annotation detection and novel protein characterization.

## 3 Using existing database annotations, local structural alignment, and LLMs to screen SAE features for cohesiveness

### 3.1 Combining protein annotation databases identifies structurally and functionally cohesive features

We expand annotation coverage by incorporating the full InterPro database (Hunter et al. [2009]), which includes annotations from UniProtKB, Pfam, CATH / Gene3D (Orengo et al. [1997]), (Buchan et al. [2002]), and 19 other sources. We evaluate associations at the protein level rather than amino acid level, allowing us to capture cases where feature activations occur near—but not exactly on—annotated elements, and enabling us to combine both protein-level annotations (like Pfam domains) and more granular annotations (like binding sites and motifs in UniProtKB). We also include hierarchical codes such as Pfam clans and CATH topologies to capture broader biological concepts.

For each SAE feature, we sample up to 1100 proteins across 10 activation levels (up to 100 proteins each for activation 0, 0.1, etc up to 1.0) and test whether any single code can predict high versus low feature activation, calculating F1 scores between predicted and actual activations. **This approach doubles our annotation coverage**. As shown in Figure 2, expanding the databases increases the percentage of features with F1 > 0.8 from approximately 25% to over 60%. The highest-performing annotations are distributed relatively evenly across UniProtKB, Pfam clans, Gene3D/CATH codes, and Pfam families, demonstrating that SAE features capture biological concepts at multiple levels of granularity. Features with moderate performance (F1 0.5-0.75) are predominantly associated with UniProtKB categories.

**Figure 1:**
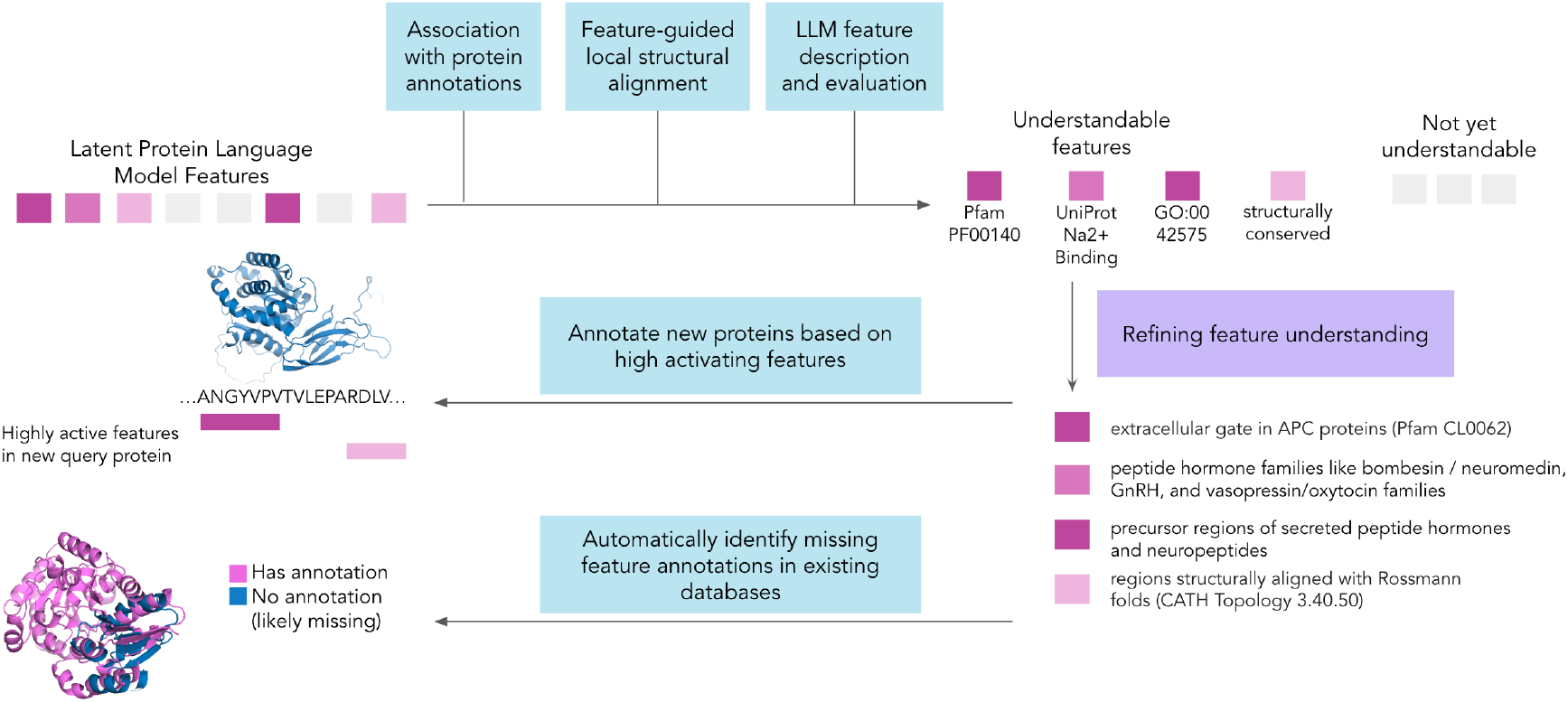
Workflow for protein latent feature applications.

**Figure 2:**
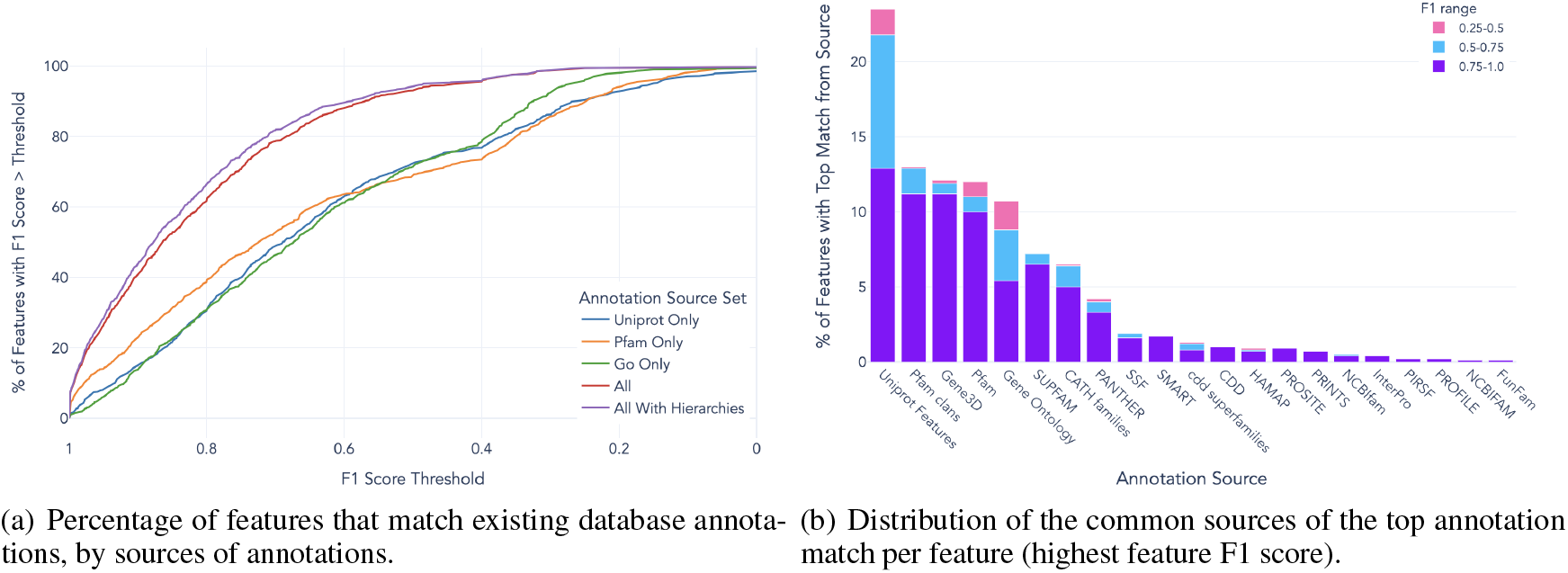
Additional databases expand our ability to find cohesive features. Hierarchical codes like Pfam clans often correspond more closely to a feature than a Pfam family, indicating the SAE learns different levels of specificity.

**This metric serves dual purposes:** quantifying annotation correspondence and screening for feature cohesiveness—features that correspond strongly to existing annotations likely represent understandable biological elements suitable for annotation purposes.

### 3.2 Feature-guided local structural alignment independently identifies structurally cohesive features

To better understand what causes these features to fire, we turn to structural analysis. This serves two benefits. First, we can track structural subdomains that may not have their own annotation. Second, we can track structural elements that fire even when an annotation is missing. InterPLM (Simon and Zou [2024]) demonstrated that when a feature associated with an existing annotation activates on a protein that lacks this corresponding annotation, what initially appears to be an erroneous feature activation can actually indicate missing or incorrect database labels. Through manual inspection, they identified three features that were correctly identifying functional patterns in proteins that should have been annotated but weren’t. However, investigating each feature individually for missing labels through manual verification would be infeasible at scale, motivating our automated pipeline for systematically identifying such annotation gaps.

To scale up this process, we introduce a procedure for Feature-Guided Local Structural Alignment to find “structurally cohesive” features, meaning the regions that activate highly have a consistent 3D structure, even if it has not been annotated. In our procedure, we sample 20 AlphaFold-predicted structures from the top activating proteins of a given SAE feature (activation above 0.7), after de-duplicating for structures from gene orthologs. We then crop selected structures to the 100 amino acids surrounding the peak activating SAE amino acid. We run pairwise local alignment between all possible pairs, and score the alignments with backbone *RMSD*_100_, a modification of Root Mean Square Deviation that is more lenient for longer alignments (Carugo and Pongor [2001]). We use this rather than RMSD because it finds longer complex conserved structures rather than, for example, a single alpha helix match. Because we are only aligning cropped structures, we can use a robust structural alignment algorithm, CEalign (Shindyalov and Bourne [1998]) with minimal cost. Additionally, by restricting alignments to regions where the SAE feature activates, we evaluate structural similarity in functionally relevant areas.

In Figure 3 we see that better structural alignment (lower *RMSD*_100_) is correlated (pearson r=0.53) with higher annotation F1 scores. Specifically, 92% of features with *RMSD*_100_ *<* 5 have a code-based F1 > .8, while only 51% of features with an *RMSD*_100_ > 5 have a code-based F1 > .8. Thus, for pairwise alignments with low *RMSD*_100_, we would expect they indeed share a local structural feature, even if one protein is missing an annotation, or is an uncharacterized novel protein. Since *RMSD*_100_ scores below 4 consistently indicate clear structural similarity, we use this as our stringent threshold. Scores between 4 and 5 also typically represent genuine structural similarity but with occasional false positives, making this our more permissive threshold for broader coverage.

**Figure 3:**
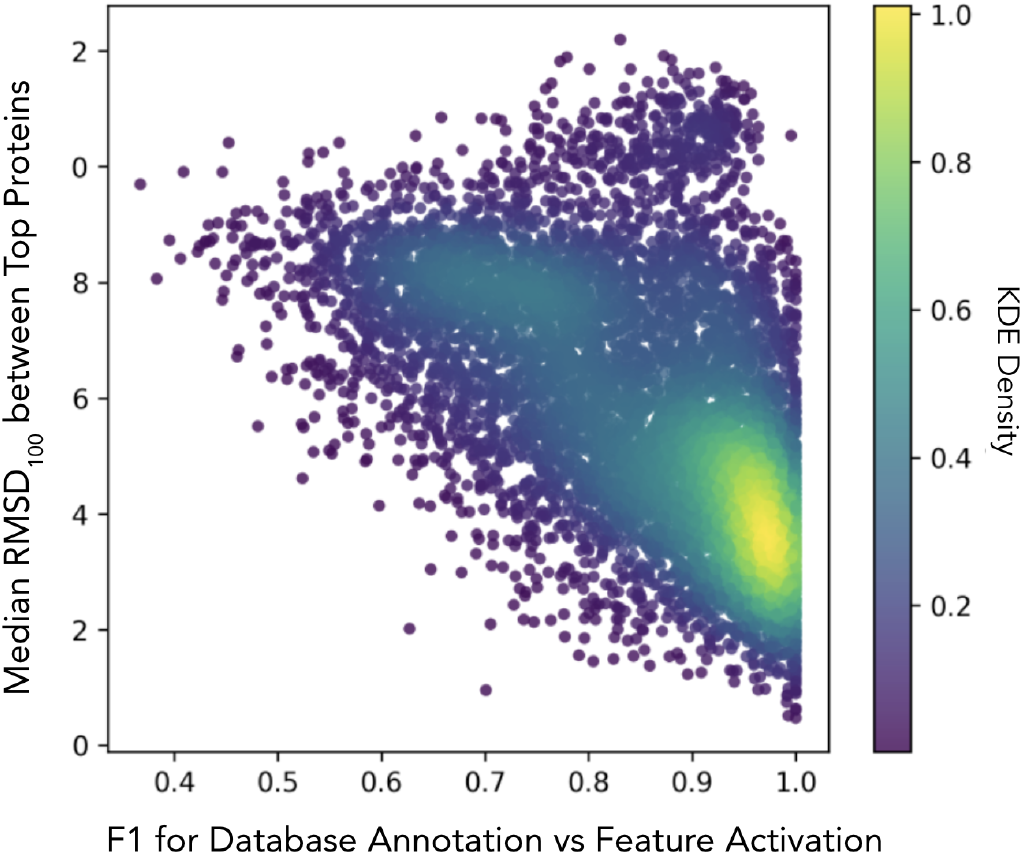
RMSD vs. F1 for existing database annotations. Many features with high *RMSD*_100_ do not correspond closely with existing annotations, while the vast majority of features with low *RMSD*_100_ correspond with existing database annotations.

### 3.3 Large Language Models identify additional cohesive features missed by other methods

While many features fire on a single database annotation code, other features appear to fire on shared traits across existing annotations. Thus, asking LLMs to reason over protein data is a natural step in developing better feature descriptions that can be used for annotation. We adapt the automated pipeline from InterPLM (Simon and Zou [2024]), using Claude-3.5 Sonnet (new) to generate feature descriptions by providing it with protein metadata from our expanded sources along with examples of 40 proteins showing varying levels of maximum feature activation. The LLM analyzes these to identify what protein and amino acid characteristics cause the feature to activate at different levels, generating descriptions of the underlying biological patterns. As validation, we test whether these LLM-generated descriptions can predict feature activation levels on held-out proteins. We evaluate the ability of the generated descriptions and the annotation metadata to classify proteins as high or low activating for each feature, calculating F1 scores on a separate test set to ensure the descriptions capture generalizable patterns rather than overfitting to the training examples.

We find that the LLM’s ability to describe a feature is highly correlated with the F1 score of the single best annotation code, as shown in Figure 4. That is, though we want the model to reason about a combination of codes, a primary driver of performance is the primary existing database annotation.

**Figure 4:**
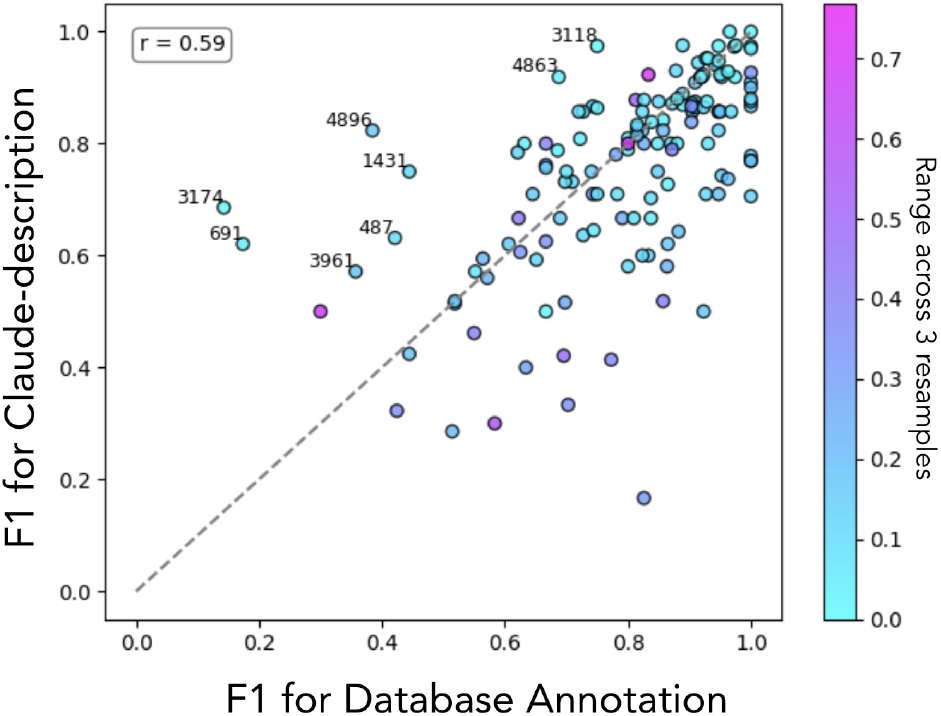
LLM-based F1 score vs. code-based F1 score on the same train/val split.

Still, we find interesting cases where the LLM quantitatively improves performance. For example, for f/3174, the LLM identifies “this feature activates on transmembrane domains in multi-pass membrane proteins, particularly those involved in protein complex assembly and ion transport across membranes.” This description, which focuses on transmembrane domains across a broader range of proteins than would be covered by a single Pfam code, allows the description to outperform existing annotation codes. Similarly, for another feature (f/4896), the LLM writes, “This feature activates on precursor regions of secreted peptide hormones and neuropeptides that undergo proteolytic processing to produce bioactive signaling molecules.” This description correctly identifies that the feature fires on several different types of peptide hormones and neuropeptides, even though they do not share a GO code.

Finally, the LLM can sometimes still interpret polysemantic features that appear to fire on two distinct protein elements. For example, f/3118 appears to fire at both the phosphohistidine of histidine kinases, and at chlorine channel sites for proton-coupled chloride transporters. While we have not been able to find a sequential or structural similarity between these two kinds of sites, the LLM correctly describes that the feature, “activates on functional domains involved in ion-mediated signaling, particularly histidine kinase domains and chloride channel domains that facilitate ion transport across membranes.” Ideally this feature would fire on a single monosemantic concept. However, this description can still reveal that the activating proteins are likely one of two types that function in a similar pathway.

## 4 Using PLM SAE features to enhance protein annotation

Having identified cohesive features using our three approaches, we now show three benefits of utilizing these features. In particular, we utilize these features to identify subdomains within existing annotations that specifically appear in the literature. Second, we utilize these features to scalably identify missing CATH annotations. Finally, we utilize these features to rapidly identify structural matches for metagenomic proteins in NMPfamsDB.

### 4.1 SAE features capture granular subdomains interpretable through targeted literature search

While our annotation matching successfully identifies coherent features, it also reveals an important limitation: we can identify the **types** of proteins where features activate, but not always what **specific elements within** those proteins cause the activation. This limitation, however, highlights a key advantage of SAE features-their ability to capture granularity beyond existing annotation schemes. Multiple SAE features often share the same top annotation code yet activate on different structural subregions, revealing subdomains that current databases treat as single units.

Figure 5 illustrates both this limitation and opportunity. Eight different features (f/253, f/515, f/1505, f/1579, f/1712, f/1731, f/2768, and f/3288) all achieve high F1 scores for the same Pfam clan annotation (CL0062), yet each activates on distinct regions–transmembrane helices, cytoplasmic domains (f/253), and extracellular domains (f/515). While our annotation-based screening identifies these as coherent features, it cannot explain their specific roles, demonstrating both the power of SAE features to decompose protein families and the need for methods to interpret this granularity.

**Figure 5:**
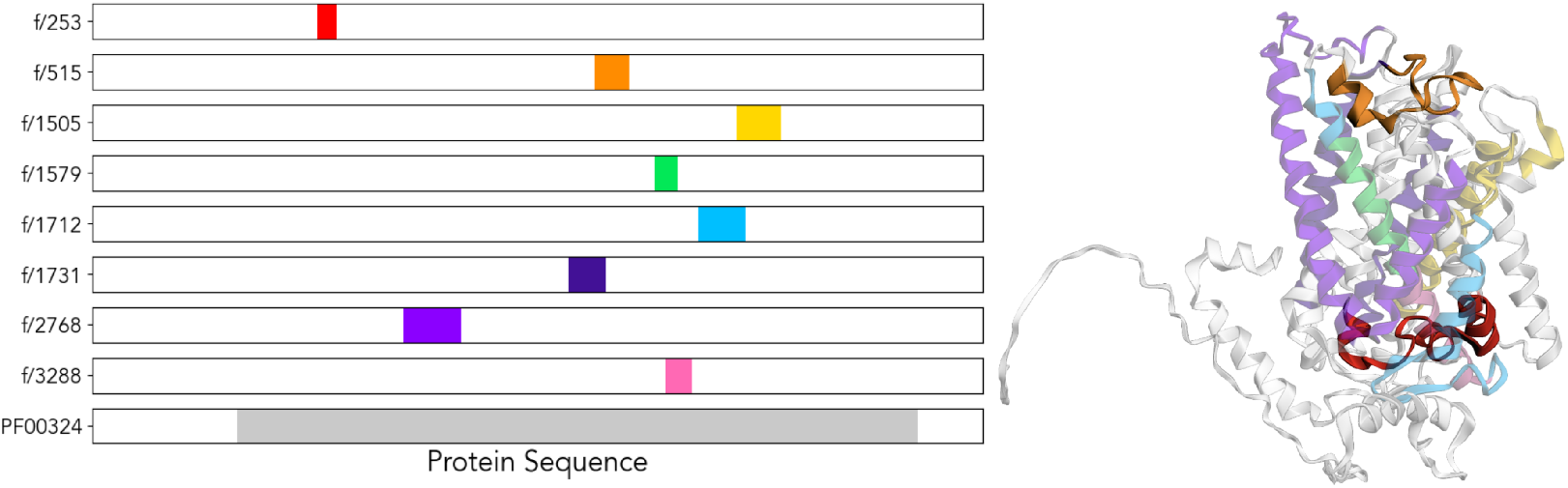
Eight SAE features correspond to the same Pfam clan (APC, CL0062) but activate on distinct structural components within GAP1_YEAST, including different transmembrane helices, cytoplasmic domains, and extracellular loops. Left: Each row shows feature activation along the protein sequence with highly activated (> 0.8) residues highlighted in color, compared to the single Pfam domain (gray). Right: Feature activations on AlphaFold predicted structure (AFDB: P19145) showing each feature highlighting distinct structural components.

To bridge this gap, we tested whether language models could retrieve literature discussing specific protein regions where features activate. We provided OpenAI’s o4-mini-high with the 10 highest-activating proteins for selected features, their gene descriptions from UniProtKB, and precise amino acid positions of peak activation, asking it to search for papers describing these regions. In a pilot of 6 features, this approach successfully identified papers that discussed those exact regions for 4 cases. For example, for f/515, the model retrieved literature on extracellular loops in APC proteins, identifying the effects of mutations in this precise region (Raba et al. [2014])—functional detail entirely absent from the broader clan annotation. An example prompt for this pilot, and more detail on the retrieved papers is in Appendix B.

This demonstrates that SAE features can reveal biologically meaningful subdomains within existing annotations, but realizing this potential requires methods to interpret their specific functional roles. While literature search shows promise for this task, it currently requires manual validation and is limited by API availability.

### 4.2 SAE features can identify missingness in existing databases

We examine proteins that activate the same SAE feature but differ in their annotation status—some have the top database annotation while others do not. When we measure *RMSD*_100_ between annotated and unannotated proteins, we often find they align just as well structurally as annotated proteins align with each other, as shown in Figure 6. This suggests the unannotated proteins may have missing database labels. For example, we note f/401, whose best code-based description is a CATH Topology (3.40.50, with F1 of .84). However, several of the top activating proteins for f/401 are not tagged with any CATH code using Gene3D. Still, these structures have very low local RMSD to structures that are tagged with the CATH code using Gene3D, suggesting the CATH code is missing.

**Figure 6:**
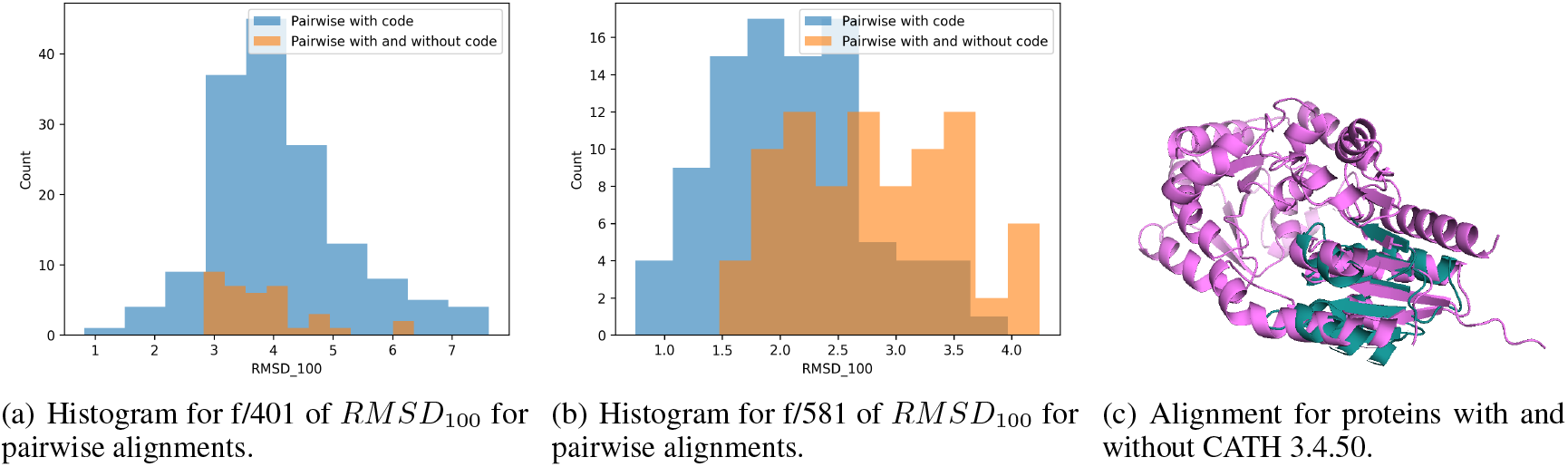
Aligning structures with the top existing annotation to structures without the annotation reveals features like (a) f/401 and (b) f/581 with strong structural similarity despite different annotations. This can identify missing annotations in existing databases, like in (c) showing an alignment for two proteins that activate on f/401 where one protein (Q0P9A8) has and one protein (Q5WSK6) is missing the CATH 3.40.50 annotation, the topology code for the Rossmann fold. For clarity, only the area around the feature is shown for Q5WSK6.

To analyze this at scale, we reviewed the 20 proteins per feature randomly sampled in Section 3.2, looking at features with an F1 of .8 or above for a CATH code or topology. Across 221 features, we find 1,055 of those proteins that are not tagged with a CATH code by Gene3D, but have local structural similarity to a protein also activating the same feature that does have a Gene3D annotation. As external validation, we compare the CATH tags from Gene3D, a sequence-based model, to TED (Lau et al. [2024]), which uses a deep learning structure-based approach. We find 491 of these proteins indeed have a hit for that same CATH topology in TED (see Appendix Table 2). For proteins that do not have hits even in TED, it is possible that these features can fire on a subdomain within a code, so the entire structure is not similar enough to be tagged.

While we can verify these seemingly missing annotations for CATH by using TED, there are SAE features that align with annotations from other databases like Pfam or UniProtKB. Thus, this combination of screening for existing annotations and local structural alignment can likely help identify potential missing annotations beyond just CATH.

### 4.3 Features can rapidly detect structural matches in unannotated metagenomic proteins

A key advantage of feature-based annotation is that granular features can detect conserved domains even when full proteins show no sequence similarity to known families. This enables annotation of highly divergent metagenomic sequences that lack Pfam matches, which we test using NMPfamsDB (Baltoumas et al. [2024]).

We find that many of our features activate highly in proteins the SAE was not trained on. Specifically, for over 50% of our features, there is at least one NMPfamsDB protein that activates that feature with a value ≥ 0.7. Then, by applying our local structural alignment procedure, we can identify 615 features with strong median local structural alignments (*RMSD*_100_ *<* 5 for pairwise alignments between one Swiss-Prot protein and one metagenomic protein) and 181 features with median *RMSD*_100_ *<* 4. This is strong evidence these features activate on the same element in both Swiss-Prot proteins and metagenomic proteins, as seen in Figure 7.

**Figure 7:**
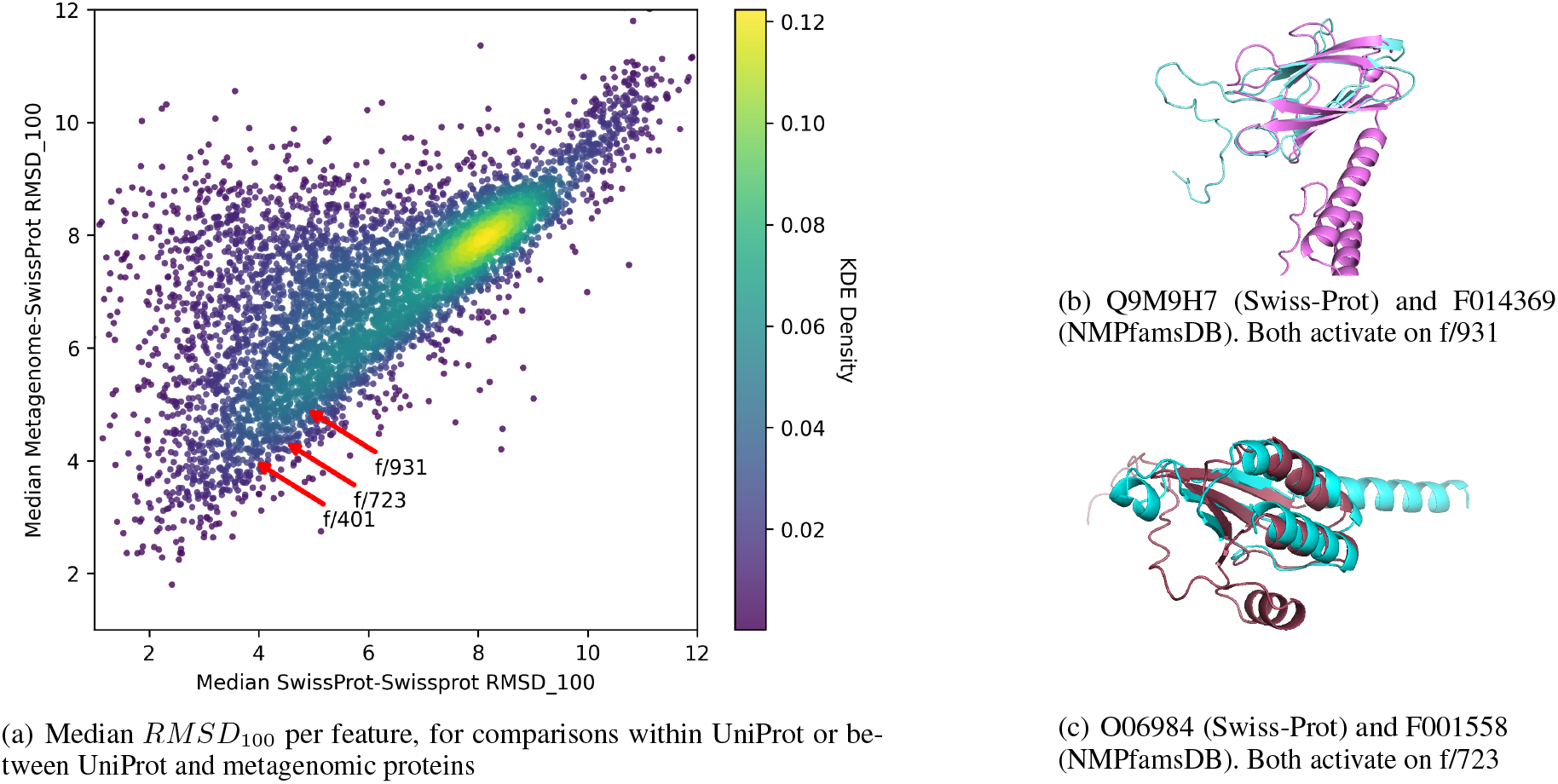
(a) *RMSD*_100_ for comparisons within Swiss-Prot or between Swiss-Prot and metagenomic proteins; (b) local structural alignment for f/931 for a Swiss-Prot (Q9M9H7) and metagenomic protein F014369; (c) local structural alignment for f/723 for a Swiss-Prot (O06984) and metagenomic protein F001558.

Using just these 615 features, we find matches between 12,526 metagenomic proteins in NMPfamsDB and Swiss-Prot proteins with an *RMSD*_100_ ≤ 5 (14.9% of the 83,878 metagenomic proteins in NMPfamsDB analyzed). At an *RMSD*_100_ threshold below 4, we find matches for 8,077 metagenomic proteins (9.6%). Many of the SAE features that align structurally in the metagenomic proteins also have high correspondence to CATH domains (378 of the 615 have a code-based F1 above 0.8 for a CATH code). Still, we also find features like f/723 and f/931 that correspond to a granular sub-domain within Pfam codes. As a final corroboration, for f/723 and f/931, we note that for several metagenomic proteins, if we run FoldSeek on the protein, several of the top Swiss-Prot hit also activate this feature. That demonstrates that these SAE features can be used to find the same structural element found by FoldSeek. However, using SAE features has a natural advantage over FoldSeek, in that we can provide the known structural or functional information about the feature(s) that triggered the alignment.

## 5 Conclusion

In this work, we demonstrate the benefits of using latent features from protein language models for annotation. We find features that consistently activate on discernible subdomains, though these cannot yet be understood automatically. We also find features that can identify missing database annotations at scale, and find features that allow us to characterize metagenomic proteins. We can pre-compute the top proteins for each feature, tie each feature to existing databases, and generate LLM-descriptions for each feature. Thus, for a new protein, this workflow returns not only a structurally similar characterized protein in constant time, but also information about exactly the aligned region.

We note several important limitations of the work: first, while LLMs can help in pulling relevant literature, identifying where within proteins these features fire remains a manual task due to hallucinations and the need for careful verification. Second, our local structural validation approach likely misses features that span distant regions or involve flexible structural motifs. Third, for now we rely on a single layer (Layer 18) of ESM-2 650M and focus our annotation validation on Swiss-Prot, which limits the space of proteins we can match.

Future work should expand analysis across multiple layers within PLMs and systematically evaluate how the quality of the latent features influences our ability to find structurally consistent matches. Additionally, more rigorous benchmarking against existing annotation approaches like FoldSeek and Merizo-search can highlight the benefits of each method. We expect advances in LLM capabilities and more advanced protein representations may improve further on these tasks, and hope this framework can provide a useful proof-of-concept as this field develops further.

## A Flexible regions

We note that many features still have a high code-based F1 even though they also have high *RMSD*_100_. Some of these are regions that are structurally flexible, for example f/73 below. We see three proteins, each with a homeodomain-like region highlight in orange and blue, and the highly activated region for f/73 in pink.

**Figure 1:**
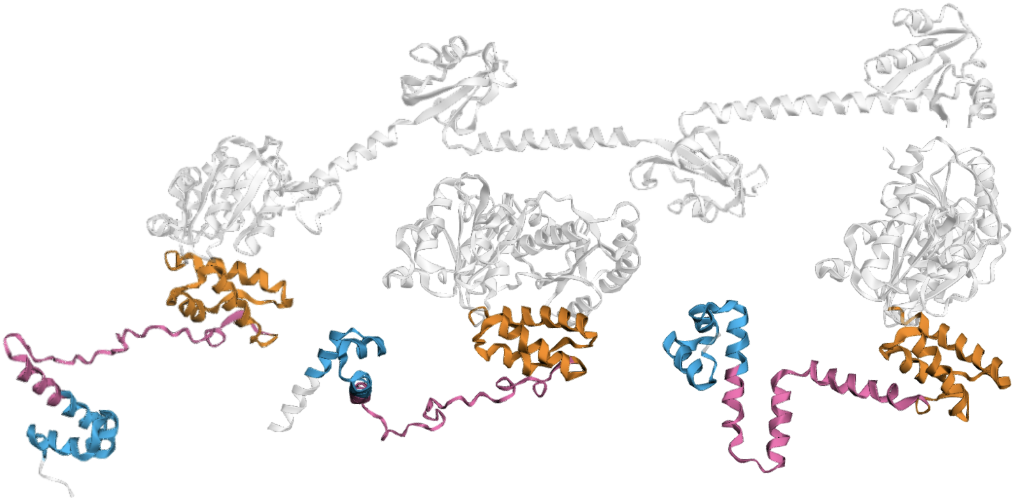
f/73 captures a variety of structures that all link two structurally consistent domains. Here, three proteins have homeolike-domains are in orange and blue, while the highly active region for f/73 is in pink

## B Understanding granular features

For six features, we used o4-mini-high to attempt to find citations that discussed the specific regions of interest. An example prompt is given below, followed by the best citation (where applicable) found by the model. Each citation returned by the model was reviewed manually, as the model could sometimes hallucinate specific quotes or mutations that were not found in the underlying papers. Where no relevant citation was retrieved, the feature was analyzed manually to determine its specific function. Sometimes very promising literature about the specific region exists, but was not returned by o4-mini-high, perhaps because of only asking about up 10 gene-species combinations per feature. For example, manually searching for additional papers revealed that f/253 corresponds to the cytoplasmic loop between transmembrane domain 2 and 3 in a topological model of amino acid permeases Cosgriff and Pittard [1997].

**Prompt for f/515**

What do we know about the following proteins in these amino acid regions mentioned for each? Cite papers that look specifically at or very near these regions

ACTP_PECCP Cation/acetate symporter ActP (Acetate permease) (Acetate transporter ActP) in Pectobacterium carotovorum subsp. carotovorum (strain PC1) at 332

MNTH_AGRFC Divalent metal cation transporter MntH in Agrobacterium fabrum (strain C58 / ATCC 33970) (Agrobacterium tumefaciens (strain C58)) at 309

…

KUP_CYTH3 Probable potassium transport system protein Kup in Cytophaga hutchinsonii (strain ATCC 33406 / DSM 1761 / CIP 103989 / NBRC 15051 / NCIMB 9469 / D465) at 271

KUP_STRA1 Probable potassium transport system protein Kup in Streptococcus agalactiae serotype Ia (strain ATCC 27591 / A909 / CDC SS700) at 277

PUTP_STAAN Sodium/proline symporter (Proline permease) in Staphylococcus aureus (strain N315) at 310 In total (across all proteins) provide me only with 2-4 of the best matching citations and what we know about the region from each paper

**Table 1:** o4-retrieved feature citations for specific sub-domains.

| Feature | o4 description of best citation | Paper link |
| --- | --- | --- |
| 73 | No citation retrieved discussed this specific subdomain (a flexible linker region sandwiched between HTH domains). | — |
| 253 | No citation retrieved discussed this specific subdomain (in the cytoplasmic loop between TM2 and TM3). | — |
| 401 | “Schmidt *et al.* solved crystal structures of the *Aquifex aeolicus* Kdo transferase (WaaA), a GT-B family homolog, revealing that the loop encompassing residues 98–102 (equivalent to *E. coli* position ~101) shapes the acceptor–substrate binding site...” | https://pubmed.ncbi.nlm.nih.gov/22474366/ |
| 515 | “Site-directed mutagenesis targeting extracellular loop 4 of *S. aureus* PutP (residues 310, 314, 318) showed that altering the amino acid at position 310 completely abolishes proline uptake...” | https://www.sciencedirect.com/science/article/pii/S0969212614000835?utm_source=chatgpt.com |
| 711 | “In vivo cysteine cross-linking between TM2 and TM8—including sites around residue 316—demonstrated that MurJ adopts both inward- and outward-facing states during transport. Disruption of membrane potential selectively destabilized the inward-facing conformation...” | https://pubmed.ncbi.nlm.nih.gov/30482840/ |
| 931 | “Residue 88 (human numbering) lies in the β-strand F of the TTR fold, forming part of a critical hydrogen-bond network that stabilizes the tetramer...” | https://pmc.ncbi.nlm.nih.gov/articles/PMC8122960/ |

## C Missing CATH annotations

Below we show the first 100 missing CATH annotations identified by our workflow, of the 491 that match with TED annotations. We considered a match if TED contained a code matching the top CATH code for a given feature that was within the same toplogy. Best code represents the top CATH code for that feature, while TED label is the exact TED label for that protein (may either be a CATH homologous superfamily or topology).

**Table 2:** Rows 1–50.

| Protein | Feature | Best code | TED label |
| --- | --- | --- | --- |
| Q01473 | 4 | 1.20.120.160 | 1.20.120 |
| Q7Y0V9 | 52 | 3.30.530.20 | 3.30.530.20 |
| Q0WV12 | 52 | 3.30.530.20 | 3.30.530.20 |
| Q9FVI6 | 52 | 3.30.530.20 | 3.30.530.20 |
| Q8XA02 | 57 | 3.10.105.10 | 3.10.105.10 |
| P46890 | 57 | 3.10.105.10 | 3.10.105.10 |
| P77172 | 81 | 3.30.70.270 | 3.30.70.270 |
| Q58121 | 112 | 1.20.58.340 | 1.20.58 |
| O28044 | 112 | 1.20.58.340 | 1.20.58 |
| Q01473 | 119 | 1.20.120.160 | 1.20.120 |
| Q49430 | 121 | 1.20.1560.10 | 1.20.1560.10 |
| Q0VFX2 | 129 | 1.20.5.500 | 1.20.5 |
| Q5BL57 | 129 | 1.20.5.500 | 1.20.5 |
| Q499U4 | 129 | 1.20.5.500 | 1.20.5 |
| Q25C79 | 129 | 1.20.5.500 | 1.20.5 |
| Q4UMJ9 | 159 | 1.20.1250.20 | 1.20.1250.20 |
| P28246 | 159 | 1.20.1250.20 | 1.20.1250.20 |
| Q9JXM5 | 169 | 1.25.40.10 | 1.25.40.10 |
| O51072 | 180 | 1.25.40.10 | 1.25.40.10 |
| Q8K4P7 | 180 | 1.25.40.10 | 1.25.40.10 |
| Q8RWN0 | 194 | 3.40.395.10 | 3.40.395.10 |
| Q0WKV8 | 194 | 3.40.395.10 | 3.40.395.10 |
| Q09275 | 194 | 3.40.395.10 | 3.40.395.10 |
| O13769 | 194 | 3.40.395.10 | 3.40.395.10 |
| Q8L7S0 | 194 | 3.40.395.10 | 3.40.395.10 |
| Q2PS26 | 194 | 3.40.395.10 | 3.40.395.10 |
| Q54KW6 | 196 | 1.25.40.10 | 1.25.40 |
| P34511 | 218 | 1.25.40.20 | 1.25.40.20 |
| P18540 | 251 | 3.40.50.2300 | 3.40.50.2300 |
| O22232 | 281 | 3.80.10.10 | 3.80.10 |
| O04615 | 310 | 2.60.210.10 | 2.60.210.10 |
| Q9FKD7 | 310 | 2.60.210.10 | 2.60.210.10 |
| Q9XHZ8 | 310 | 2.60.210.10 | 2.60.210.10 |
| Q8C008 | 371 | 2.60.40.10 | 2.60.40 |
| Q9STL8 | 399 | 3.40.140.10 | 3.40.140.10 |
| Q9FG71 | 399 | 3.40.140.10 | 3.40.140.10 |
| O82264 | 399 | 3.40.140.10 | 3.40.140.10 |
| Q9LYC2 | 399 | 3.40.140.10 | 3.40.140.10 |
| Q04368 | 399 | 3.40.140.10 | 3.40.140.10 |
| P76349 | 401 | 3.40.50.2000 | 3.40.50.2000 |
| D3DJ42 | 401 | 3.40.50.2000 | 3.40.50.20 |
| Q5WSK6 | 401 | 3.40.50.2000 | 3.40.50.2000 |
| Q9SH31 | 419 | 3.40.50.2000 | 3.40.50 |
| Q3E9A4 | 419 | 3.40.50.2000 | 3.40.50.2000 |
| Q9LFP3 | 434 | 3.40.50.2000 | 3.40.50.2000 |
| P75207 | 442 | 1.20.1560.10 | 1.20.1560.10 |
| Q49430 | 443 | 1.20.1560.10 | 1.20.1560.10 |
| Q9ZD06 | 449 | 3.30.2350.10 | 3.30.2350.10 |
| Q8FB47 | 449 | 3.30.2350.10 | 3.30.2350.10 |
| Q8ZGM2 | 449 | 3.30.2350.10 | 3.30.2350.10 |

**Table 3:** Rows 51–100.

| Protein | Feature | Best code | TED label |
| --- | --- | --- | --- |
| Q92HG4 | 449 | 3.30.2350.10 | 3.30.2350.10 |
| P77607 | 544 | 3.40.50.1390 | 3.40.50.1390 |
| Q8XA02 | 607 | 3.10.105.10 | 3.10.105.10 |
| P46890 | 607 | 3.10.105.10 | 3.10.105.10 |
| A9NCA7 | 713 | 3.40.50.2000 | 3.40.50.2000 |
| O67214 | 713 | 3.40.50.2000 | 3.40.50.2000 |
| P37597 | 751 | 1.20.1250.20 | 1.20.1250.20 |
| O04292 | 789 | 3.30.450.20 | 3.30.450.20 |
| Q9Z7F1 | 789 | 3.30.450.20 | 3.30.450.20 |
| A2XBL9 | 789 | 3.30.450.20 | 3.30.450.20 |
| Q39123 | 789 | 3.30.450.20 | 3.30.450.20 |
| B0K165 | 843 | 3.30.479.30 | 3.30.479.30 |
| P32233 | 848 | 3.40.50.300 | 3.40.50.300 |
| Q2NL82 | 848 | 3.40.50.300 | 3.40.50.300 |
| A0A0H2URH2 | 849 | 3.40.50.2000 | 3.40.50.2000 |
| A0A0H2URJ6 | 849 | 3.40.50.2000 | 3.40.50.2000 |
| P33694 | 849 | 3.40.50.2000 | 3.40.50.2000 |
| A1JSF2 | 873 | 3.40.50.300 | 3.40.50.300 |
| P50837 | 889 | 3.30.420.10 | 3.30.420.10 |
| Q60953 | 889 | 3.30.420.10 | 3.30.420.10 |
| P53296 | 891 | 3.30.559.10 | 3.30.559.30 |
| I1S097 | 894 | 3.90.550.10 | 3.90.550.10 |
| D3ZZN9 | 936 | 2.60.40.150 | 2.60.40.150 |
| Q9C8E6 | 936 | 2.60.40.150 | 2.60.40.150 |
| P54739 | 972 | 3.30.200.20 | 3.30.200.20 |
| P54735 | 972 | 3.30.200.20 | 3.30.200.20 |
| C4JDF8 | 1003 | 1.10.1200.10 | 1.10.1200.10 |
| P39404 | 1013 | 3.40.50.2300 | 3.40.50.2300 |
| O83933 | 1016 | 1.20.1600.10 | 1.20.1600 |
| P63400 | 1039 | 1.10.1760.20 | 1.10.1760 |
| O67248 | 1039 | 1.10.1760.20 | 1.10.1760 |
| Q58299 | 1053 | 3.60.40.10 | 3.60.40.10 |
| O29259 | 1119 | 1.10.443.10 | 1.10.443.10 |
| P07261 | 1119 | 1.10.443.10 | 1.10.443.20 |
| O83202 | 1205 | 3.30.70.270 | 3.30.70.270 |
| P77172 | 1205 | 3.30.70.270 | 3.30.70.270 |
| Q2NKC0 | 1205 | 3.30.70.270 | 3.30.70.270 |
| Q10419 | 1233 | 2.40.30.170 | 2.40.30.170 |
| P55501 | 1239 | 3.30.420.10 | 3.30.420.10 |
| Q44493 | 1320 | 2.150.10.10 | 2.150.10.10 |
| P75800 | 1329 | 3.30.70.1230 | 3.30.70.270 |
| B5XZP2 | 1347 | 1.20.1250.20 | 1.20.1250.20 |
| D0CCT2 | 1347 | 1.20.1250.20 | 1.20.1250.20 |
| E0T2N0 | 1347 | 1.20.1250.20 | 1.20.1250.20 |
| O34353 | 1360 | 2.120.10.80 | 2.120.10.30 |
| Q10412 | 1380 | 1.20.920.10 | 1.20.920 |
| Q6YRK2 | 1416 | 3.30.70.270 | 3.30.70.270 |
| B0XZV4 | 1447 | 1.20.1250.20 | 1.20.1250.20 |
| O83837 | 1466 | 1.25.40.10 | 1.25.40 |
| Q8IAR5 | 1482 | 2.30.42.10 | 2.30.42.10 |

## References

Etowah Adams, Liam Bai, Minji Lee, Yiyang Yu, and Mohammed AlQuraishi. From mechanistic interpretability to mechanistic biology: Training, evaluating, and interpreting sparse autoencoders on protein language models. bioRxiv, pages 2025–02, 2025.

Fotis A Baltoumas, Evangelos Karatzas, Sirui Liu, Sergey Ovchinnikov, Yorgos Sofianatos, I-Min Chen, Nikos C Kyrpides, and Georgios A Pavlopoulos. Nmpfamsdb: a database of novel protein families from microbial metagenomes and metatranscriptomes. Nucleic Acids Research, 52(D1):D502–D512, 2024.

Alex Bateman, Lachlan Coin, Richard Durbin, Robert D Finn, Volker Hollich, Sam Griffiths-Jones, Ajay Khanna, Mhairi Marshall, Simon Moxon, Erik LL Sonnhammer, et al. The pfam protein families database. Nucleic acids research, 32(suppl_1):D138–D141, 2004.

Daniel WA Buchan, Adrian J Shepherd, David Lee, Frances MG Pearl, Stuart CG Rison, Janet M Thornton, and Christine A Orengo. Gene3d: structural assignment for whole genes and genomes using the cath domain structure database. Genome research, 12(3): 503–514, 2002.

Oliviero Carugo and Sándor Pongor. A normalized root-mean-spuare distance for comparing protein three-dimensional structures. Protein science, 10(7): 1470–1473, 2001.

UniProt Consortium. Uniprot: a hub for protein information. Nucleic acids research, 43(D1):D204–D212, 2015.

Angela J Cosgriff and AJ Pittard. A topological model for the general aromatic amino acid permease, arop, of escherichia coli. Journal of bacteriology, 179(10): 3317–3323, 1997.

Sarah Hunter, Rolf Apweiler, Teresa K Attwood, Amos Bairoch, Alex Bateman, David Binns, Peer Bork, Ujjwal Das, Louise Daugherty, Lauranne Duquenne, et al. Interpro: the integrative protein signature database. Nucleic acids research, 37(suppl_1):D211–D215, 2009.

Philip Jones, David Binns, Hsin-Yu Chang, Matthew Fraser, Weizhong Li, Craig McAnulla, Hamish McWilliam, John Maslen, Alex Mitchell, Gift Nuka, et al. Interproscan 5: genome-scale protein function classification. Bioinformatics, 30(9): 1236–1240, 2014.

John Jumper, Richard Evans, Alexander Pritzel, Tim Green, Michael Figurnov, Olaf Ronneberger, Kathryn Tunyasuvunakool, Russ Bates, Augustin Žídek, Anna Potapenko, et al. Highly accurate protein structure prediction with alphafold. nature, 596(7873): 583–589, 2021.

Shaun M Kandathil, Andy M Lau, Daniel WA Buchan, and David T Jones. Foldclass and merizo-search: Scalable structural similarity search for single-and multi-domain proteins using geometric learning. Bioinformatics, page btaf277, 2025.

Eli Levy Karin and Martin Steinegger. Cutting edge deep-learning based tools for metagenomic research. National Science Review, page nwaf056, 2025.

Andy M Lau, Shaun M Kandathil, and David T Jones. Merizo: a rapid and accurate protein domain segmentation method using invariant point attention. Nature Communications, 14(1): 8445, 2023.

Andy M Lau, Nicola Bordin, Shaun M Kandathil, Ian Sillitoe, Vaishali P Waman, Jude Wells, Christine A Orengo, and David T Jones. Exploring structural diversity across the protein universe with the encyclopedia of domains. Science, 386(6721):eadq4946, 2024.

Zeming Lin, Halil Akin, Roshan Rao, Brian Hie, Zhongkai Zhu, Wenting Lu, Nikita Smetanin, Robert Verkuil, Ori Kabeli, Yaniv Shmueli, et al. Evolutionary-scale prediction of atomic-level protein structure with a language model. Science, 379(6637): 1123–1130, 2023.

Christine A Orengo, Alex D Michie, Susan Jones, David T Jones, Mark B Swindells, and Janet M Thornton. Cath–a hierarchic classification of protein domain structures. Structure, 5(8): 1093–1109, 1997.

Michael Raba, Sabrina Dunkel, Daniel Hilger, Kamila Lipiszko, Yevhen Polyhach, Gunnar Jeschke, Susanne Bracher, Johann P Klare, Matthias Quick, Heinrich Jung, et al. Extracellular loop 4 of the proline transporter putp controls the periplasmic entrance to ligand binding sites. Structure, 22(5): 769–780, 2014.

Ilya N Shindyalov and Philip E Bourne. Protein structure alignment by incremental combinatorial extension (ce) of the optimal path. Protein engineering, 11(9): 739–747, 1998.

Elana Simon and James Zou. Interplm: Discovering interpretable features in protein language models via sparse autoencoders. bioRxiv, pages 2024–11, 2024.

Michel van Kempen, Stephanie S Kim, Charlotte Tumescheit, Milot Mirdita, Cameron LM Gilchrist, Johannes Söding, and Martin Steinegger. Foldseek: fast and accurate protein structure search. Biorxiv, pages 2022–02, 2022.

Yang Zhang and Jeffrey Skolnick. Tm-align: a protein structure alignment algorithm based on the tm-score. Nucleic acids research, 33(7): 2302–2309, 2005.

